# Automated real-time EEG sleep spindle detection for brain state-dependent brain stimulation

**DOI:** 10.1101/2022.06.05.494865

**Authors:** Umair Hassan, Gordon Feld, Til Ole Bergmann

## Abstract

Sleep spindles are a hallmark electroencephalographic (EEG) feature of non-rapid eye movement (NREM) sleep and believed to be instrumental for sleep-dependent memory reactivation and consolidation. However, direct proof of their causal relevance is hard to obtain, and our understanding of their immediate neurophysiological consequences is limited. To investigate their causal role, spindles need to be targeted in real-time with sensory or non-invasive brain stimulation techniques. While fully automated offline detection algorithms are well established, spindle detection in real time is highly challenging due to their spontaneous and transient nature. Here, we present the *real-time spindle detector* (RTSD), a robust multi-channel EEG signal processing algorithm that enables the automated triggering of stimulation during sleep spindles in a phase-specific manner. We validated the RTSD method by streaming pre-recorded sleep EEG datasets to a real-time computer system running a Simulink® Real-Time™ implementation of the algorithm. Sleep spindles were detected with high levels of sensitivity (∼83%) and precision (∼78%) and an F1-score of ∼0.81 in reference to state-of-the-art offline algorithms (which reached similar levels when compared to each other), for both naps and full nights, and largely independent of sleep scoring information. Detected spindles were comparable in frequency, duration, amplitude, and symmetry, and showed the typical time-frequency characteristics as well as a centroparietal topography. Spindles were detected close to their center and reliably at the predefined target phase. The RTSD algorithm therefore empowers researchers to target spindles during human sleep and apply the stimulation method and experimental paradigm of their choice.

## 1 Introduction

Thalamocortical sleep spindles, i.e., 0.5-2 s bursts of oscillatory brain activity at sigma frequency (∼12-15 Hz) with waxing and waning amplitude, are a hallmark feature of the electroencephalogram (EEG) during non-rapid eye movement (NREM) sleep. Spindles subserve sleep-dependent memory reactivation (Bergmann et al., 2012a) and systems consolidation (Diekelmann and Born, 2010) via their nested phase-amplitude coupling with neocortical slow oscillations (SO; 0.5-1 Hz), and hippocampal ripples (>80 Hz) (Staresina et al., 2015), supporting phase-dependent plasticity (Bergmann & Born, 2018) and the synaptic rescaling of cortical neurons (Klinzing et al., 2019). To investigate their neurophysiological underpinnings and test their causal role for memory consolidation, spindles need to be targeted and manipulated experimentally in humans using sensory or transcranial brain stimulation in a brain state-dependent manner (Bergmann 2018), as done successfully for the SO.

SO phase-dependent transcranial magnetic stimulation (TMS) of the left human primary motor cortex (M1) during NREM sleep revealed that corticospinal excitability, as assessed by motor evoked potentials (MEPs), were smaller during SO down-than up-states (Bergmann et al., 2012b). Using closed-loop SO-phase triggered auditory stimulation, Ngo et al. (2013, 2015) were able to enhance SO amplitude, associated spindle generation, and memory consolidation. And targeted memory reactivation (TMR) studies successfully explored the benefit of targeting reactivation cues phase-locked to SO up- or down-states (Göldi et al., 2019; Shimizu et al. 2018).

While SOs are an easy real-time target, also faster oscillations can be targeted using modern real-time systems (Zrenner et al., 2018). During wakefulness, real-time EEG-triggered TMS of M1, demonstrated phase-dependent MEP modulations during the sensorimotor mu-alpha rhythm (Bergmann et al., 2019; Zrenner et al., 2018), and TMS bursts repeatedly targeting the more excitable mu-alpha troughs induced long-term potentiation (LTP)-like MEP increases (Zrenner et al., 2018). While alpha amplitude modulations are slower and more easily predictable than spontaneous spindle bursts of the sigma band, spindles have a better signal-to-noise ratio.

Several automated offline spindle detection approaches are available (Vallat and Walker, 2021; Lacourse et al., 2019; Weber, 2013; Warby et al., 2014; Parekh et al., 2017), including machine or deep learning-based detection methods (Kaulen et al., 2022; LaRocco et al., 2018; Kulkarni et al., 2019) with a potential for real-time applications, but actual implementations are scarce. Lustenberger et al. (2016) targeted ongoing spindles with transcranial alternating current stimulation (TACS) but provided limited information regarding the real-time algorithm and its phase specificity. The *Portiloop* system, a portable system on chip, at least achieved real-time spindle detection with an F1-Score of 0.71 by training artificial neural networks on a field-programmable gate array (FPGA) (Valenchon et al., 2021).

Here we present the *real-time spindle detector (RTSD)*, a robust, empirically validated, and fully automated real-time signal processing algorithm that is able to identify spindles from ongoing EEG recordings close to their amplitude maximum (spindle center) and send out a phase-specific trigger signal. We validated the RTSD using real-time streaming of pre-recorded sleep data from naps and full nights, comparing its performance to several state-of-the-art offline spindle detection algorithms, and evaluating key characteristics of the detected spindles.

## 2 Materials and Methods

### 2.1 The real-time spindle detector

The real-time spindle detector (RTSD) identifies 0.5 to 2 s long distinct oscillatory patterns of waxing and waning amplitude in the sigma band (12-15 Hz) in ongoing EEG data. Unlike offline algorithms that can rely on long data segments comprising complete spindles, the RTSD has to work in a causal manner, i.e., by analyzing only the present and past data points. To robustly detect spindles while they unfold and therefore with incomplete information about their final shape, duration, and amplitude, the RTSD relies on the analysis of several complementary high-level signals derived from the raw data, which are evaluated in parallel and combined to make a decision. Also, personalized detection criteria related to individual spindle power and peak frequency cannot be taken from the full data of the same sleep recording but have to be derived from baseline sleep data recorded either during the beginning of the same sleep session or during a previous sleep session of the same subject (e.g., an adaptation night). In the latter case, spindle peak frequency and root mean square (RMS) of the spindle band power can be extracted using established offline detection procedures that utilize envelope-based detection, such as YASA, A7, or SpiSOP (Lacourse et al., 2019; Vallat and Walker, 2021; Weber, 2013).

#### 2.1.1 Hardware and software Implementation

The RTSD algorithm is implemented in a real-time Simulink (MathWorks) environment in MATLAB r2017b, which can be compiled, loaded, and run on any Simulink compatible real-time computer system. For actual real-time detection, digitized EEG/PSG data are streamed via a real-time compatible EEG amplifier with digital output streaming (e.g., NeurOne Tesla EEG system, Bittium, Finland or ActiChamp Plus, BrainProducts, Germany). For validation purposes in the present work, we replayed existing datasets to the compiled real-time Simulink model running on a dedicated real-time system (*bossdevice*, sync2brain Germany). The BEST toolbox [23] was used to control the Simulink model, setting individual thresholds for spindle detection on the bossdevice. Raw time series data of the detected events, such as spindles and SOs as well as the instantaneous EEG phases at the time of detection, were recorded in data files for confirmatory post-hoc offline analysis. In our implementation, the RTSD algorithm had a loop delay of 5-15 ms, which is induced primarily by the hardware, the interface software (here, Simulink), and the digital bandpass filters.

#### 2.1.2 Data preprocessing

Raw data are processed through a number of custom spatial filters, i.e., a linear combination of the streamed channels, resulting in an arbitrary number of virtual channels for which spindles could be detected independently in parallel. Every 10 ms, 520 ms of the most recent data are extracted and used for further analyses, resulting in a 98% overlap of consecutively extracted analysis windows. Then, by applying a two-pass (zero-phase) least-squares finite impulse response (FIR) filter of order 20 to the most recent 520 ms data segment we create two fundamental signals: using a passband of 1-30 Hz a broadband EEG signal (EEG_bb)_ is constructed, and using a passband of the individual spindle frequency ± 2 Hz a sigma band EEG signal (EEG_σ_) is created. Then, 10 ms from each side of EEG_bb_ and EEG_σ_ are removed to create 500 ms segments of data that are free of filter edge artifacts. The filtered EEG_bb_ and EEG_σ_ signals are then further processed to derive four high-level signals, which are described in the following section together with their respective thresholds.

#### 2.1.3 Computation of high-level signals

##### EEG_σ_ RMS power signal

To gain an index of absolute spindle power, the root mean square (RMS) is computed using a 250 ms moving data window of EEG_σ_ with a step size of 100 ms to construct the EEG_σ_ RMS power signal. The RMS power threshold is individualized for each subject based on their baseline sleep data by taking the mean of the EEG_σ_ RMS power plus 1.5 times their standard deviation.

##### EEG_σ_ Relative power signal

To prevent broadband power changes from being mistaken for spindle-related increases in the sigma band, the EEG_**σ**_ RMS power signal needs to be normalized. A fast Fourier transform of the EEG_bb_ is used to calculate the EEG_σ_ relative power. A 500 ms moving data window is zero padded on both sides (250 ms each) to 1 s to construct a frequency spectrum with 1 Hz resolution. The power summed across the frequency bins corresponding to the spindle peak frequency ± 2 Hz (i.e., the individual sigma band) is divided by the power summed across the 1-30 Hz bins to construct the EEG_σ_ Relative power signal. An increase in EEG_σ_ Relative power is caused by a local increase in the power of the sigma band but not by broadband power increases that occur proportionally across the entire spectrum. A fixed threshold value of 15-20% relative power is generally sufficient to estimate sigma band specific activity, but the threshold can also be determined individually based on the baseline sleep data.

##### EEG_σ_ x EEG_bb_ Correlation signal

To prevent artifacts or noise in the raw data to be mistaken for actual spindle events in the sigma bandpass filtered data, the actual waveform shape of the potential spindle is evaluated in the raw data. For this purpose, the Pearson correlation coefficient between EEG_bb_ and EEG_σ_ is calculated for a 250 ms moving data window. A high correlation coefficient indicates that the bandpass filtered data reflects actual activity of the sigma band rather than artifacts or noise. A fixed threshold value of r ≥ 0.65 is generally sufficient to detect N2 and N3 spindles, but the value may need to be adopted based on the signal-to-noise ratio (SNR) of spindles in the subject and the specific spatial filters applied.

##### EEG_bb_ Instantaneous frequency signal

To further ensure the other high-level signals are not confounded by activity in neighboring frequency bins (which for offline detection methods can be ensured by the use of higher-order bandpass filters), only spindle oscillations within certain frequency bounds are accepted. Therefore, the instantaneous frequency is calculated from the time derivative of the Hilbert phase of the EEG_bb_ signal over a 250 ms moving data window. Then, it is determined for which percentage of the data window the EEG_bb_ instantaneous frequency remains within the range of the individual spindle peak frequency ± 5x the standard deviation of the spindle frequency (Hz). A fixed threshold value of 75% generally indicates a stable instantaneous frequency in the sigma band.

#### 2.1.4 Spindle detection

Eventually, a spindle is detected when at least three out of the four above-described high-level signals satisfy their respective criteria and in addition the spindle duration criterion is met. This is the case when time since the EEG_σ_ RMS power signal crossed the ‘entry threshold’ of mean + 1.15 standard deviations is greater than 250 ms, which corresponds to half of the minimum spindle duration of 500 ms, allowing for the identification of the spindle center also for very short spindles. Maximum spindle power (i.e., the assumed spindle center) is detected when the time derivative of the EEG_σ_ RMS power crosses zero and the RMS power begins to descend. The end of the spindle is detected once the signal falls below the ‘entry threshold’ again. Spindle detection is suspended when the RMS power remains above the ‘entry threshold’ for more than 2 s, and it resumes only after at least three of the four signals again exceed their respective thresholds.

#### 2.1.5 Real-time spindle phase estimation

In addition to detecting spindle events, also their oscillatory phase is estimated in real-time using the *phastimate* algorithm (Zrenner et al., 2020). In summary, the phase of the spatially filtered raw signal is continuously estimated by band-pass filtering the most recent 256 ms moving data window by a least-squares FIR filter with an order of 65 and a passband of the individual spindle frequency ± 2 Hz. Then, 32.5 ms on each side of the data segment are removed to create a data segment free of filter edge artifacts. The resulting 191 ms long data segment is then used for 65 ms forward prediction using a Yule-Walker 15^th^ order autoregressive model (McFarland and Wolpaw, 2008; Chen et al., 2013). The forward predicted 65 ms data segment provides ± 32.5 ms around time point zero. A fixed time offset equal to the software and hardware delays is then applied to the zero time point to identify the actual time point zero (i.e., “now”). A static range test is applied to the phase at this time point “now” obtained from the Hilbert phase time series of the forecasted signal to identify peaks, troughs, rising flanks, and falling flanks. Phases estimated during the detected spindle event are considered spindle phases.

### 2.2 Empirical validation of the real-time spindle detector

#### 2.2.1 Sleep EEG/PSG datasets

We validated our real-time algorithm in two independent datasets of EEG and polysomnography (PSG) recordings from nocturnal naps (Dataset 1) and full nights (Dataset 2), respectively. *Dataset 1* consisted of N = 20 subjects (12 females, mean age: 23 ± 5 years) recorded during the stimulation-free adaptation session of an unpublished EEG-triggered TMS experiment conducted at the Neuroimaging Center (NIC) of the Johannes Gutenberg University Medical Center, Mainz. Subjects had no history of neurological or psychiatric disease, were right-handed, and did not take any medications. They were not allowed to drink alcohol or caffeine on the day of the experiment, and they had to be awake for at least 8 hours before the experiment to ensure moderate sleep pressure. Experimental procedures conformed to the Declaration of Helsinki and were approved by the Ethics Committee of the Landesärztekammer Rheinland-Pfalz. EEG and PSG were recorded using a 64-channel EEG-cap with sintered Ag/AgCl electrodes (Multitrodes, EasyCap) containing the following electrodes according to 10-20 EEG system: Fp1, Fp2, Fpz, AF7, AF3, AFz, AF4, AF8, F7, F5, F3, F1, Fz, F2, F4, F6, F8, FT9, FT7, FC5, FC3, FC1, FC2, FC4, FC6, FT8, FT10, T7, C5, C3, C1, Cz, C2, C4, C6, T8, TP7, CP5, CP3, CP1, CPz, CP2, CP4, CP6, TP8, P7, P5, P3, P1, Pz, P2, P4, P6, P8, PO7, PO3, POz, PO4, PO8, O1, Oz, O2, M1 (left mastoid), M2 (right mastoid); Reference, FCz; Ground, POz. For PSG, electromyography (EMG) at the chin and the vertical and horizontal electro-oculogram (VEOG and HEOG) were recorded using bipolar electrode montages. EEG and PSG data were digitized in DC mode with a 1250 Hz anti-aliasing low-pass filter and 5 kHz sampling rate using a TMS-compatible 24-bit amplifier (NeurOne Tesla with Digital-Out Option, Bittium) connected to an 8V battery. The subjects slept in a soundproof and electromagnetically shielded sleeping cabin (Desone, Germany) on a comfortable mattress resting on a wooden bed in a horizontal position. They were covered with a blanket and their heads were stabilized by a vacuum cushion. *Dataset 2* consisted of N = 10 subjects selected from the placebo session of a previously published study [17] that examined the effects of blocking metabotropic glutamate receptor 5 on sleep-dependent memory consolidation (Feld et al., 2021). The data were collected at the sleep lab of the Institute of Medical Psychology and Behavioral Neurobiology, University of Tübingen, Germany. EEG and PSG were recorded using Ag/AgCl electrodes containing the following electrodes according to 10-20 EEG system: F4, Fz, F4, C3, Cz, C4, P3, Pz, P4; Referenced to linked mastoids; Ground electrode on the forehead. For PSG, EMG at the chin and VEOG and HEOG were recorded using bipolar electrode montages. EEG and PSG data were digitized in DC mode with an 80 Hz low-pass filter and a sampling rate of 250 Hz. Further details regarding the sleep characteristics of all subjects are available in **Supplementary Tables S1 and S2**.

#### 2.2.2 Offline spindle detection

We selected three previously published and frequently used offline spindle detection algorithms for comparison and validation of our RTSD method, namely YASA [9], A7 [10], and SpiSOP [11]. These offline algorithms are actively maintained as open-source software contributions, are well documented, and easy to use. Since the A7 spindle detector itself had been shown to outperform five previously published offline detection methods [18-22], we considered no further methods than the three mentioned ones for comparison. To ensure fair comparisons, the key spindle detection criteria for both our RTSD algorithm as well as all offline spindle detection algorithms were kept identical wherever possible, namely, to detect spindles in the 12-15 Hz frequency band, with a minimum duration of 0.5 and maximum duration of 2 s, and an amplitude of mean + 1.5 times the standard deviation of the RMS bandpass filtered signal. We used published software implementations of YASA (version 0.6.01) in Python release 3.10 and of A7 (version 1.1.2) and SpiSOP in MATLAB r2017b.

For comparison with our real-time spindle algorithm, we selected channel C3 from Dataset 1 (nap recordings) re-referenced against the average of both mastoids. For Dataset 2 (full night recordings), we selected channel C3, already referenced against a linked mastoids during the original recordings. Additionally, EMG, HEOG and VEOG from PSG were selected and used in the automatic sleep scoring implemented in YASA. To separate the effect of the different spindle detection algorithms from the potential impact of the sleep scoring, spindle detection was performed twice, once on the entire data irrespective of sleep stage, and once restricted to sleep stages N2 and N3 as obtained from YASA automated sleep stage scoring. YASA scoring results were used for all validation runs with offline algorithms as well as the RTSD algorithm. For the naps of Dataset 1, a manual check of the automatic sleep stage scoring was performed for quality control as recommended for naps in the YASA software.

#### 2.2.3 Performance assessment

We assessed the performance of the RTSD against each of the offline spindle detectors, using the respective offline algorithm as ground truth. Specifically, we assessed the number of correct spindle detections (True Positive, TP), of incorrect spindle detections (False Positive, FP), and of missed spindles (False Negative, FN). Correct rejections (True Negatives, TN) cannot reasonably be assessed for rare events in continuous data and are therefore not provided. Based on these numbers, we determined Sensitivity (i.e., the percentage of detected spindles that are true spindles), Precision (i.e., the percentage of true spindles that are detected spindles), and the F1-Score (an established compound index for assessing detection algorithms combining sensitivity and precision) as defined below:

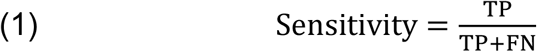

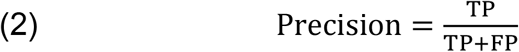

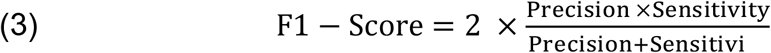

The RTSD was compared to YASA, A7, and SpiSOP. As a benchmark for the maximally expected levels of Sensitivity, Precision, and F1-Score, all three state-of-the-art offline detectors were also tested against each other, resulting in a total of six different comparisons (Real-time vs YASA, Real-time vs A7, Real-time vs SpiSOP, YASA vs A7, YASA vs SpiSOP, and A7 vs SpiSOP). Using the *method of intersection* (Lacourse et al., 2019; LaRocco et al., 2018; Warby et al., 2014), the ratio between the intersection and union of the respectively detected spindle intervals was calculated for each spindle, and a spindle match (TP) was determined whenever the ratio was greater than 0.2. Unmatched spindles found in the offline detectors were categorized as FN, and unmatched spindles found in the real-time detector were categorized as FP.

To further characterize the spindles detected in real-time, (i) morphological properties of the detected spindles, such as spindle frequency, amplitude, duration, and symmetry of wave shape, were calculated and compared with offline detected spindles as described by Purcell et al., (2017), (ii) time-frequency representations (TFR) were calculated to verify their typical narrow band waxing and waning pattern, and (iii) topographical representations of spindle power were plotted to verify their distinct localization with respect to the EEG montage selected for detection.

Time-frequency analysis of spindle related oscillatory power changes was carried out using the FieldTrip toolbox (Oostenveld et al., 2011). For all spindles, raw spatially filtered data segments of -1 to 1 s around the real-time detected spindle center were retrieved from the real-time system (bossdevice). Data were then preprocessed by de-meaning and a 1 Hz high pass filter. Time-frequency representations (TFR) were calculated separately for True Positive and False Positive spindles (compared against YASA) by using a Hanning taper windowed FFT (time steps: 50 ms, frequency steps: 0.5 Hz). TFRs were plotted as percentage change in power from the baseline at -1 to 1 s. Sigma power topographies were plotted for the time interval of -500 to 500 ms as the percentage change in power from the baseline at -550 to -500 ms.

To test whether spindle phase targeting worked as expected, spindle phase was additionally validated with the phase-detection module enabled and 4 different spindle phase targets (peak, falling flank, trough, and rising flank) in 4 additional detection runs: Post-hoc we determined the actual phase of the trigger signal based on the Hilbert transform of the sigma bandpass filtered (individual sigma peak ± 2Hz) EEG signal and evaluated the phase targeting precision using the circular standard deviation method of CircStat toolbox (Berens, 2009).

## 3 Results

### 3.1 Performance of RTSD is comparable to offline algorithms

When limiting spindle detection to NREM epochs the following performance indices were obtained. The grand average **F1-score** for RTSD vs. the different offline algorithms was 78-84% (compared to 80-81% between the offline algorithms) for the naps, and 79-83% (compared to 81% between the offline algorithms) for the full-nights. The grand average **Sensitivity** for RTSD vs. the different offline algorithms was 80-87% (compared to 83-85% between the offline algorithms) for the naps, and 81-86% (compared to 83-86% between the offline algorithms) for the full-nights. The grand average **Precision** for RTSD vs. the different offline algorithms was 76-80% (compared to 77-78% between the offline algorithms) for the naps, and 78-81% (compared to 78-80% between the offline algorithms) for the full-nights. See **Table 1** for all grand average values and **Figure 1** for distributions and subject-wise performance indices. When spindles were identified without limiting the search of the algorithms to NREM sleep stages, Sensitivity, Precision, and F1-scores were overall slightly reduced for both naps and full night recordings but still better for comparisons between RTSD and offline detection than between the offline detectors themselves when providing no sleep staging information (**Supplementary Figure S1** and **Table 1**).

**Table 1.**
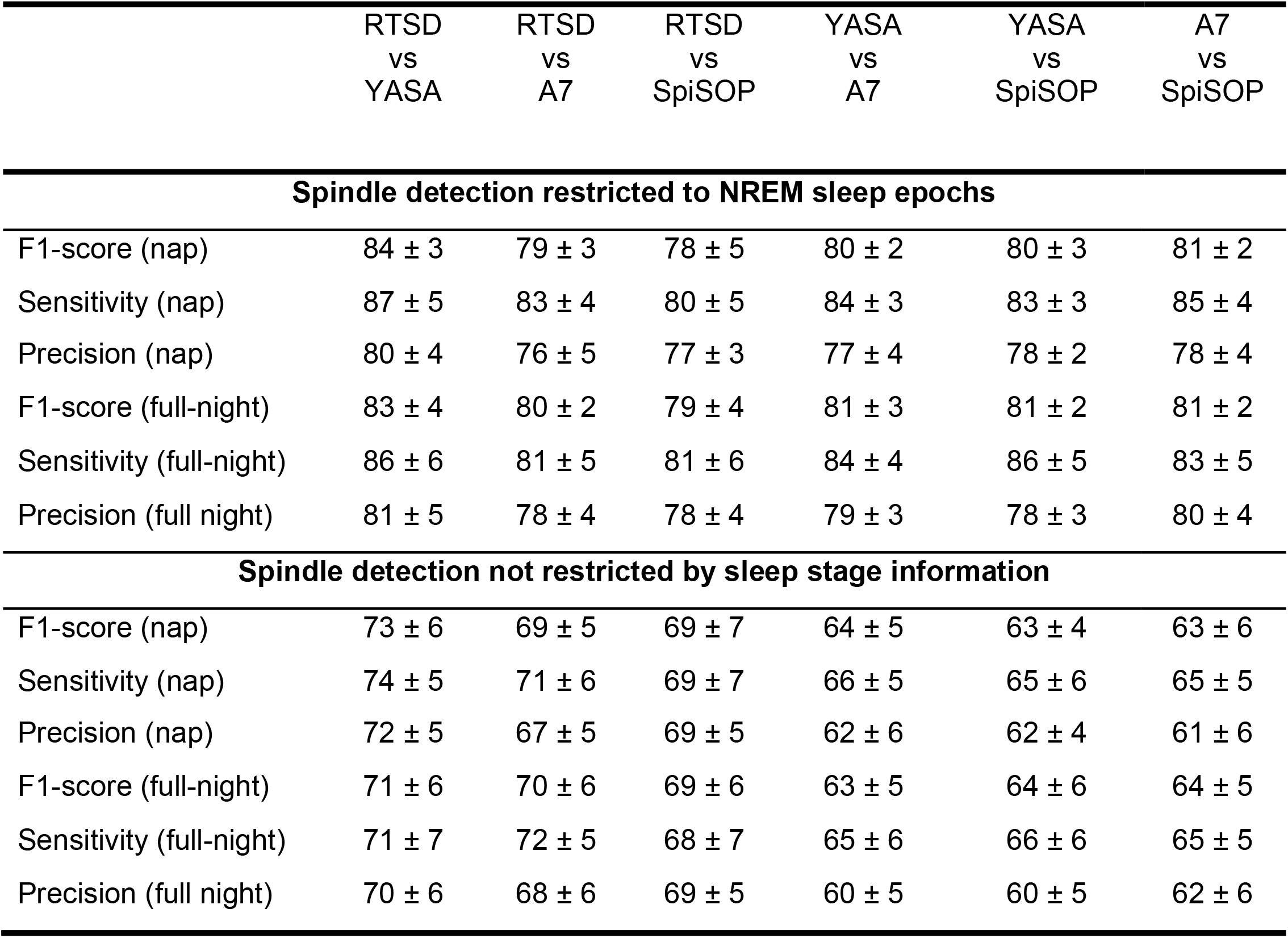
Grand average values of performance characteristics in percent (%). Means (± Standard deviation) are reported.

**Figure 1:**
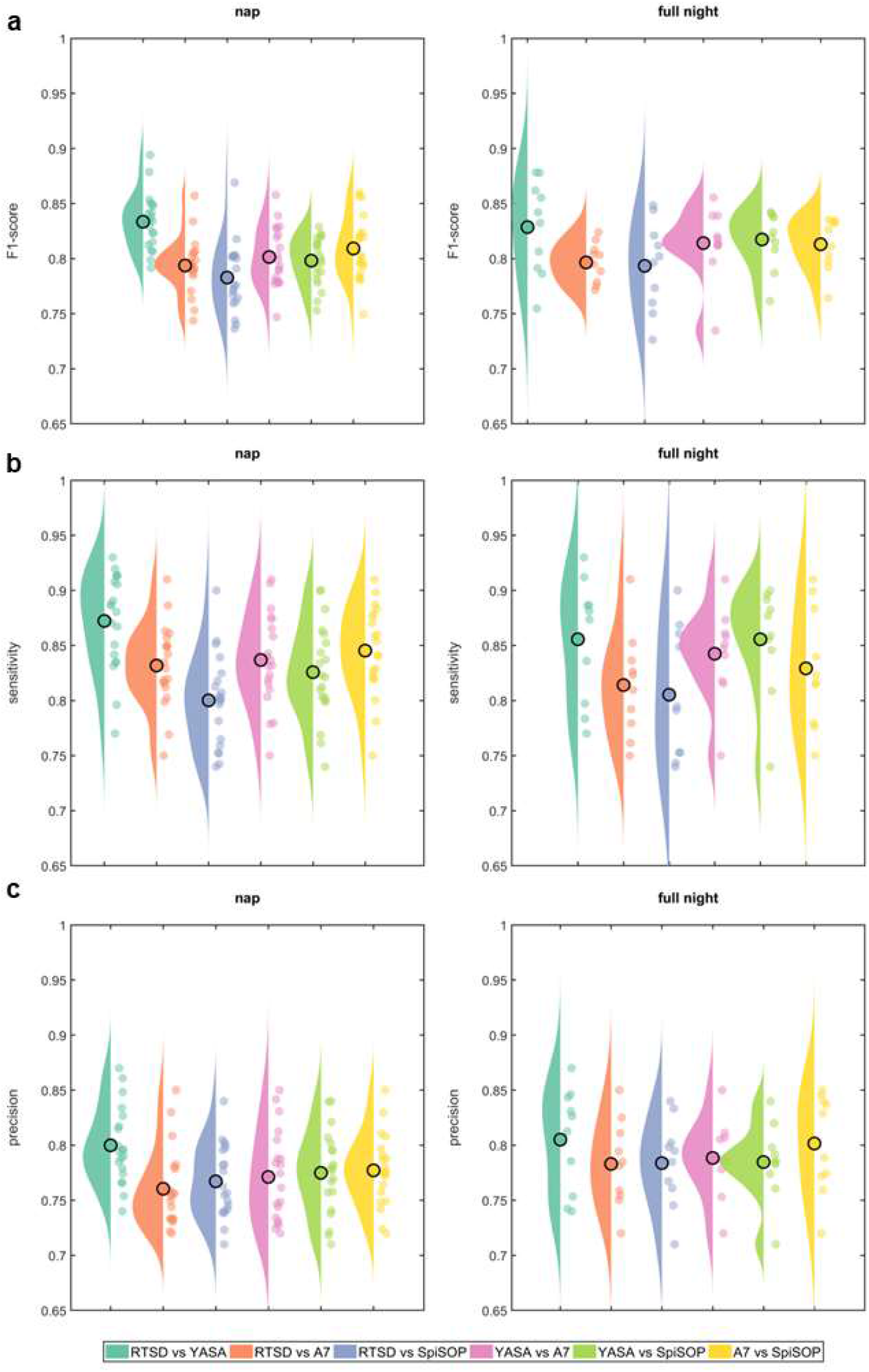
Performance values for comparisons of RTSD and offline algorithms including NREM sleep epochs only. RTSD was compared against offline algorithms (YASA, A7, and SpiSOP) and in between offline algorithms for the nap recordings (dataset 1) and full-night recordings (dataset 2) for ***A***, F1-scores, ***B***, Sensitivity, and ***C***, Precision. Single -subject datapoints (colored filled circles) and raincloud plots are provided in addition to the condition mean (black open circle)

### 3.2 Properties of real-time detected spindles are comparable to those detected offline

When comparing the distributions of spindle frequency, amplitude, duration, and symmetry of the spindle amplitude envelope for the events detected by the RTSD and the different offline algorithms (confined to NREM sleep epochs), highly similar distributions were obtained (**Figure 2**), confirming the high contingency levels described above and no systematic differences in the type of spindles being detected.

**Figure 2:**
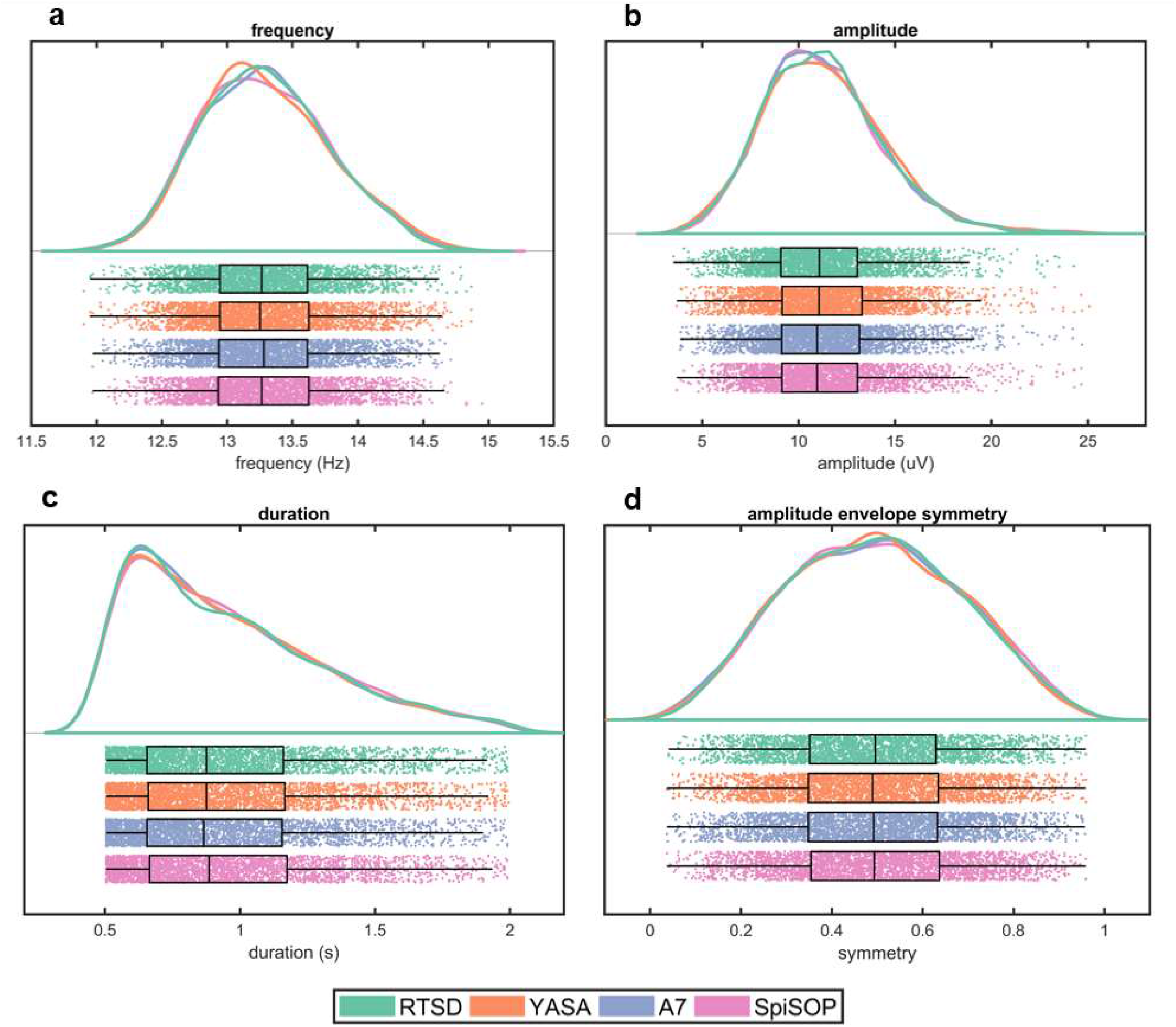
Morphological properties of real-time and offline detected spindles. Distributions across all detected spindles of all subjects and data points of all individual spindles events are provided for morphological spindle traits extracted by the same procedure, being highly comparable with respect to ***A***, frequency (Hz). ***B***, amplitude (μV). ***C***, duration (s), and ***D***, symmetry index of the spindle amplitude envelope.

Time-frequency representations (**Figure 3A**) and topographical distribution of sigma power (**Figure 3B**) of true positive spindles detected using RTSD revealed that the true positive spindles (here defined in comparison to YASA) were detected as intended and that neither adjacent frequencies nor oscillatory activity from other sources had confounded the RTSD procedure.

**Figure 3:**
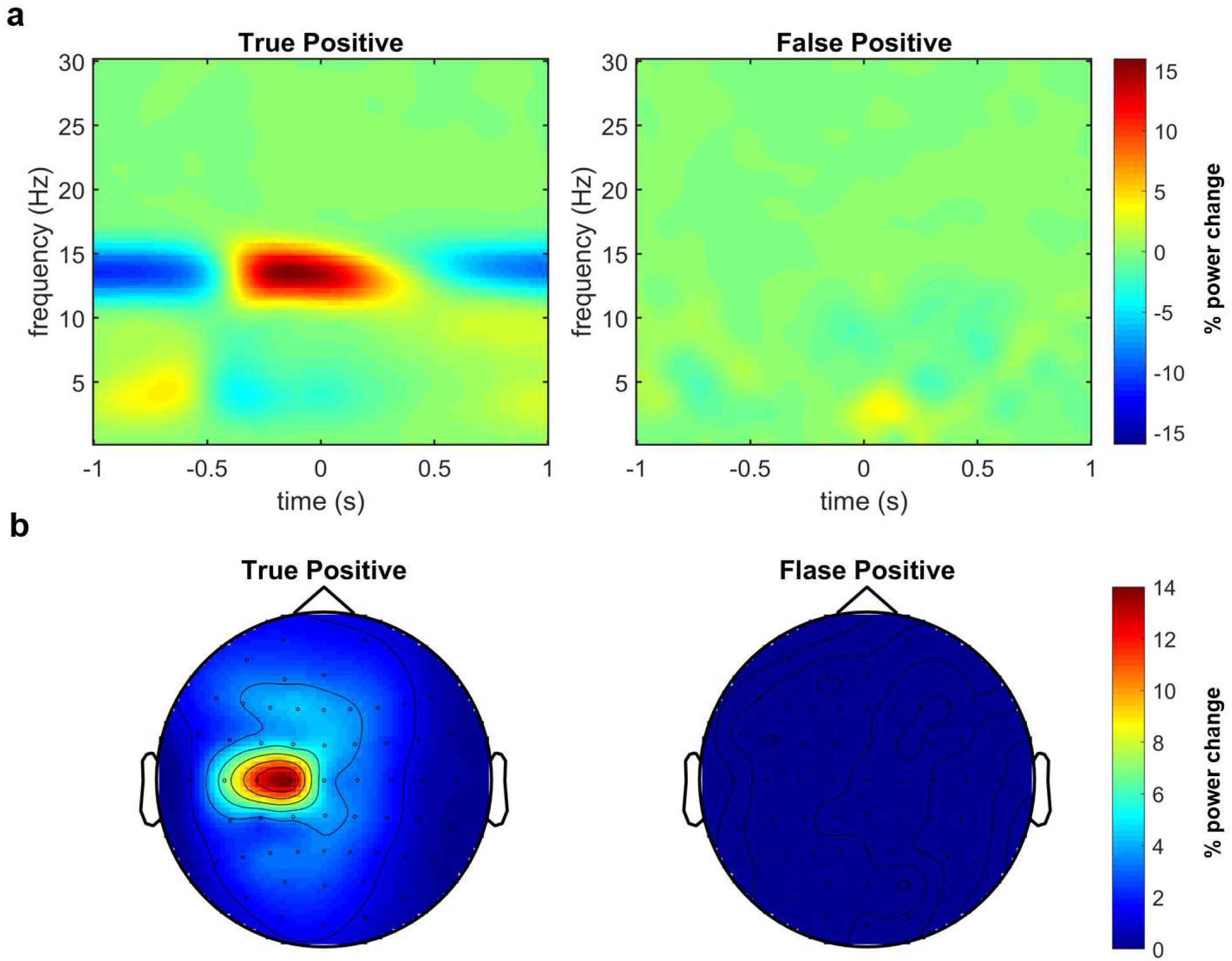
Topographical distribution and time-frequency representation of real-time detected spindles. ***A***, Time-frequency representations (TFRs) of oscillatory power in the C3 signal re-referenced against linked mastoids, calculated separately for RTSD detected True Positive and RTSD False Positive spindles (as compared to YASA) with a baseline interval of -1 to 1s. TFRs of True Positive spindles show a modulation of sigma power over time, with a relative increase from -500 to 500 ms around spindle center and relative decrease before -500 and after 500 ms. False positive spindles showed no particular pattern in the TFR that would be indicative of a particular temporal or frequency source of EEG signals leading to false alarms. ***B***, Topographical maps of the sigma power modulation from -500 to 500 ms (taken from ***A***) for True Positive and False Positive spindles detected by RTSD with a baseline interval of -550 to -500 ms. The topography of True Positive spindles verifies a local sigma power increase over C1 to C3 (specific to the target montage), suggesting that also local spindles can be targeted using RTSD. False positive spindles showed no particular topography that would be indicative of a particular spatial source of EEG signals leading to false alarms.

### 3.3 Spindle phase targeting was reliable

To verify that the instantaneous oscillatory phase of an ongoing spindle can be reliably determined in real-time, we confirmed the phase targeting post-hoc by extracting the phase information from the Hilbert transform of the individually sigma bandpass filtered signal. We did this separately for the four target phases (peak, trough, rising flank, falling flank), but collapsed across both Datasets while including only periods of NREM sleep (**Figure 4**). All spindle phases were correctly targeted with an average circular standard deviation of 53.4^°^ across all phase angles (peak: 53.9^°^; falling flank: 55.1^°^; trough: 54.6^°^; rising flank: 50.2^°^).

**Figure 4:**
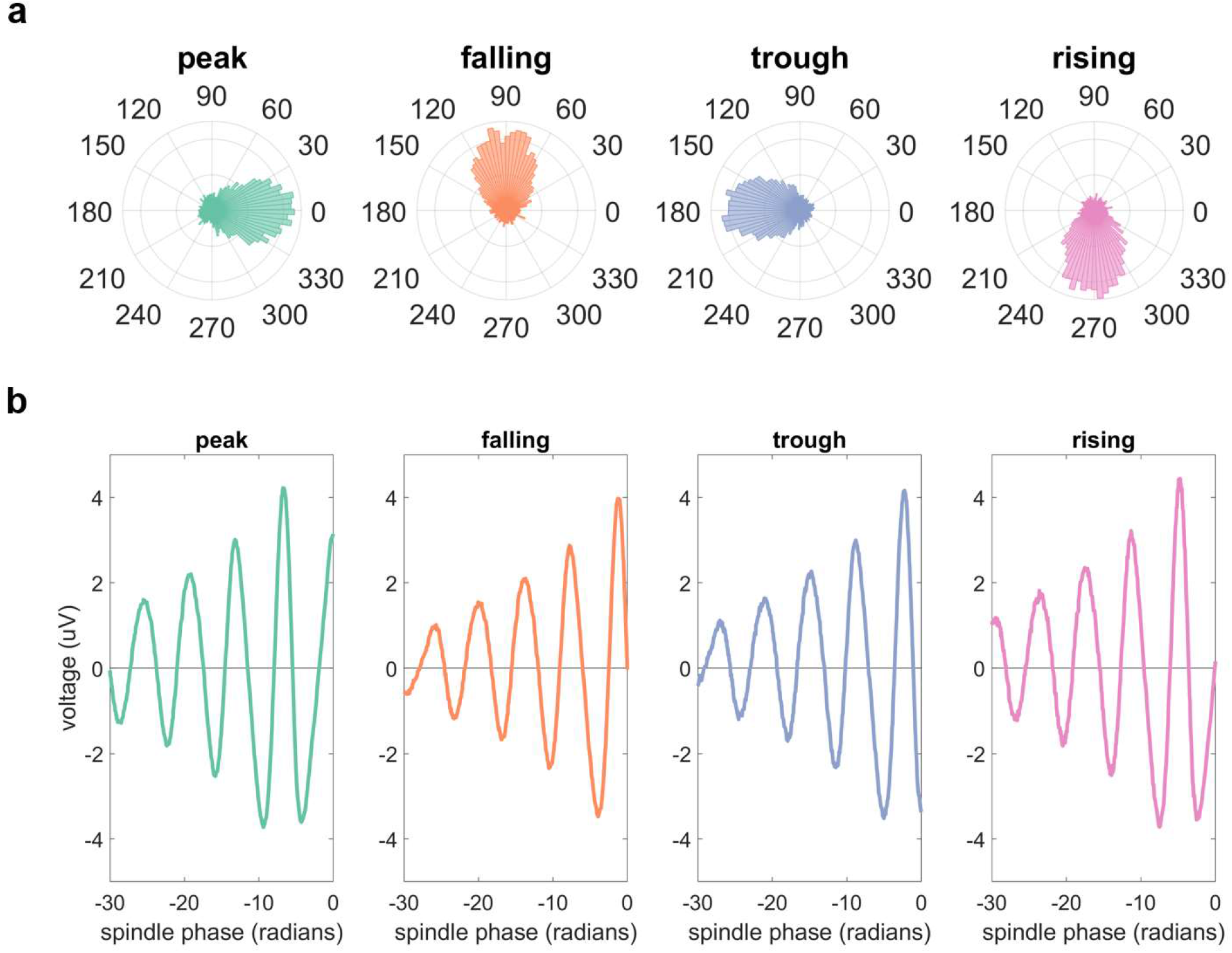
Phase estimates for different phase targets during spindles. ***A***, Phase histograms of all spindles that were detected time-locked to each of the four explicit phase targets (peak, falling flank, trough, rising flank). Actual phase angles were determined post-hoc using the Hilbert transform. ***B***, Time-locked C3 signal re-referenced against linked mastoids with the time axis transformed to phase angles (radians) of the individual spindle peak frequency before averaging across subjects to account for inter-individual differences in frequency. Spindle phase detection was successfully time locked to peaks, falling flanks, troughs, and rising flanks.

## 4 Discussion

We have presented and validated a novel method for robustly detecting sleep spindles automatically in real-time. The real-time spindle detector (RTSD) identifies ongoing spindle activity by transforming the raw EEG in several high-level signals which are evaluated in parallel and then detecting spindle activity in one or more EEG channels in a manner similar to offline spindle detection.

The performance of the RTSD was very good (F1-score around 0.8) with well-balanced sensitivity and precision levels. Higher levels were not to be expected, given that the automatic offline detection algorithms themselves did not reach higher levels when compared to each other. The F1-scores reported for other offline spindle detection methods are even slightly lower, e.g., A7 scored at 0.72 (Lacourse et al., 2019), DETOKS at 0.67 (Parekh et al., 2015), Spindler at 0.67 (LaRocco et al., 2018) and Spinky at 0.74 (Lajnef et al., 2017). However, the lower F1-scores may also be caused by the fact that these offline methods used expert labelled data as ground truth, whereas we used automatic offline spindle detectors. In addition, these methods detected spindles as short as 300 milliseconds rather than 500 milliseconds, which is slightly easier to detect by automated detection techniques. We deliberately used multiple established state-of-the-art offline spindle detection algorithms as ground truths instead of manual expert ratings, because the RTSD algorithm is not meant to compete with clinical sleep scoring procedures but to provide a novel method for neuroscientific research into the function of sleep spindles using sensory and transcranial brain stimulation approaches, for which the approximation of human judgements is less important than the use of objective and reproducible criteria. If desired, changing RTSD detection criteria allows for a flexible adjustment of sensitivity versus precision, to be as liberal or as conservative as required by the experimental design and research question.

Performance indicators were reduced only slightly for RTSD (by ∼10%) when spindle detection was not restricted to NREM sleep stages, while larger drops were observed in the consistency between offline algorithms (by ∼15-20%) (**Table 1** and **Supplementary Figure S1**). This shows that while sleep stage information improves the F1-score, the RTSD is very robust and can also be used in real-time detection settings where no sleep scoring information is available during the ongoing recordings. This robustness is remarkable, given that the RTSD has to work on less information (data) than the offline algorithms. A7, YASA, and SpiSOP require data segments of at least a few seconds to identify complete spindles, whereas the RTSD method only uses the most recent 500 ms and per definition only the first half of the ongoing spindle for detection. However, the RTSD compensates the lack of temporal information by the parallel assessment of several high-level signals derived from the incoming raw data. For example, none of the offline methods assesses whether the instantaneous frequency is within the desired bounds, which constitutes one of the four key thresholds in RTSD.

The primary limitation of the RTSD is that it locates the spindle center by detecting zero crossings of the amplitude envelope derivative as soon as the envelope begins to descend, while in reality spindle amplitude envelopes can have more than one maximum. The RTSD only detects local maxima following 250 ms of spindle onset (i.e., since the envelope has crossed the ‘entry threshold’), regardless of whether this is a global maxima. Together with the loop delay of 5-15 ms reached in our implementation (Simulink® Real-Time™ on a bossdevice from sync2brain), spindles cannot be targeted during their initial waxing phase. Removing the criterion of detecting a spindle center may allow earlier targets within a spindle but will also increase the false positive rate; whether or not this is acceptable eventually depends on the study goals. This limitation may also be overcome in the future by using additional information from the data (such as power envelope gradients) or by training deep learning approaches to predict spindles during very early phases or even before their initiation (Valenchon et al., 2021).

Using the RTSD default threshold settings, the detected spindles expressed morphological traits very similar to those detected by the offline algorithms. Moreover, time-frequency representations of the true positive spindles confirmed the typical pattern of a time limited narrow sigma band frequency increase during the spindle preceded and followed by relative decreases, which may reflect the influence of a phase-amplitude modulating SO. False positive spindles, were not associated with any particular time-frequency pattern, indicating that there was no particular systematic temporal or frequency source of EEG signals leading to false alarms. Topographical sigma power maps of true positive spindles showed the typical central topography of sleep spindles, but with a lateralization towards the left hemisphere (peaking at channel C1 and extending to C3), which is presumably owed to the left-lateralized montage (C3 vs. linked mastoids) used for spindle detection in our validation runs. Again, false positive spindles did not show a systematic sigma power distribution, indicating that no particular spatial source of EEG signals led to false alarms. Together, these auxiliary analyses confirm that the RTSD algorithm can effectively detect true sleep spindles that are very comparable to those detected by offline algorithms, without a systematic source of false alarms.

Finally, the RTSD method revealed robust real-time phase targeting (relying on the established *phastimate* algorithm; Zrenner et al., 2020), with levels of phase precision (53.4^0^ circular standard deviation) similar to comparable targets in the mu-alpha oscillation (Baur et al., 2020)). Importantly, phase targeting did not differ between isolated spindles and those occurring during a SO, showing that the overlapping asymmetrical SO/K-complex waveform with its sharp hyperpolarizing down-state did not cause any systematic bias in the spindle phase detection. This is relevant for researchers interested in studying the particular relevance of combined SO/spindle events for synaptic rescaling and memory consolidation (Klinzing et al., 2019; Latchoumane et al., 2017).

## 5 Conclusion

We have proposed and validated the *real time spindle detector* (RTSD) an automated real-time sleep spindle detection method working on single or multiple channel montages in parallel to robustly identify sleep spindles in digitally streamed EEG data. A comparison between RTSD and several state-of-the-art offline spindle detection algorithms using data from both naps and full nights of sleep revealed that RTSD performed at least as well as the offline methods, even in the absence of sleep stage information. The RTSD method therefore holds significant potential for use with real-time EEG-triggered sensory or transcranial brain stimulation to study sleep-dependent memory consolidation or synaptic plasticity in humans.

## Acknowledgments

This work was supported by funding from the Boehringer Ingelheim Foundation (BIF) and the German Research Foundation (DFG Grant 362546008) to T.O.B.

## Competing financial interests

U.H. is head of software development at sync2brain GmbH, Germany. Other authors declare no competing financial interests. The other authors declare no conflict of interest.

## Author contributions

U.H. and T.O.B. conceptualized the study. G.F. provided validation data. U.H., designed, developed, and tested the algorithm; U.H. and T.O.B wrote the manuscript; all authors jointly discussed and revised the manuscript.

## SUPPLEMENTARY MATERIALS

**Supplementary Figure S1:**
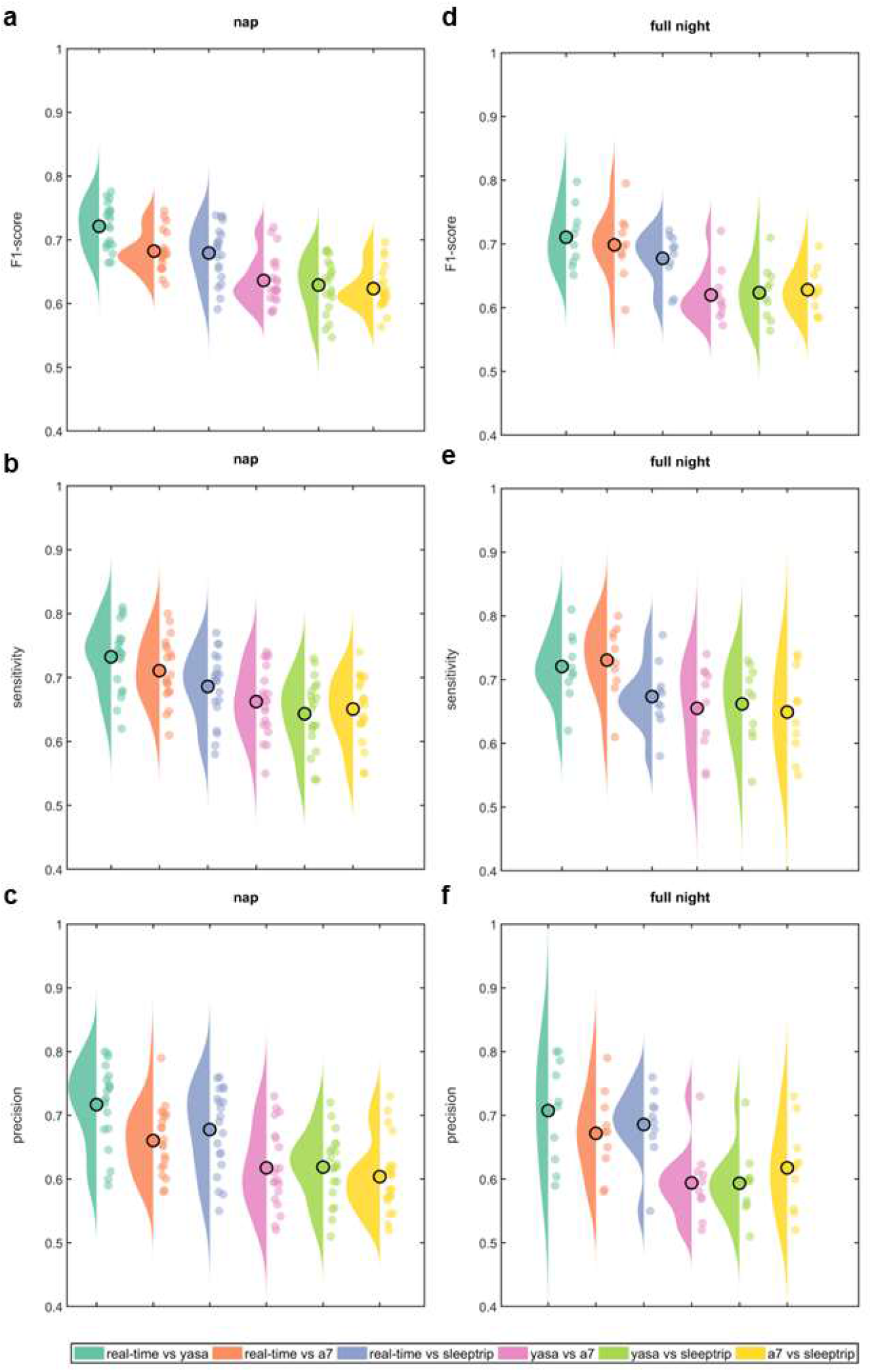
Performance values for comparisons of RTSD and offline algorithms including all sleep epochs. RTSD was compared against offline algorithms (YASA, A7, and SpiSOP) and in between offline algorithms for the nap recordings (Dataset 1) and full-night recordings (Dataset 2) for ***A***, F1-scores, ***B***, Sensitivity, and ***C***, Precision. Single -subject datapoints (colored filled circles) and raincloud plots are provided in addition to the condition mean (black open circle)

**Supplementary Table S1:** Polysomnographic sleep parameters. TST, total sleep time; N1, N2 and N3, NREM sleep stages 1, 2 and 3; SOL, sleep onset latency. Percentages are given with reference to TST. Means (± SEM) are reported.

| Parameter | Dataset 1 (nap) | Dataset 2 (full-night) |
| --- | --- | --- |
| TST | 53.5 ± 8.1 min | 426.0 ± 9.5 min |
| Wake | 7.0 ± 6.5 min | 39.7 ± 9.02 min |
| N1 | 22.1 ± 6.6 % | 11.1 ± 2.5 % |
| N2 | 50.6 ± 7.3 % | 40.9 ± 2.3 % |
| N3 | 0.95 ± 4.4 % | 23.3 ± 1.5 % |
| NREM | 82.3 ± 6.7 % | 75.3 ± 1.5 % |
| REM | 17.6 ± 6.1 % | 24.6 ± 1.4 % |
| SOL | 10.0 ± 2.8 min | 11.0 ± 2.9 min |

**Supplementary Table S2:** Number of total spindles and Mean sleep spindle density (number per minute) across C3-linked mastoids) for Dataset 1 (nap recordings) and Dataset 2 (full-night recordings) during NREM sleep.

| Subject No. | Dataset 1 Spindle number | Dataset 1 Spindle density | Dataset 2 Spindle number | Dataset 2 Spindle density |
| --- | --- | --- | --- | --- |
| 01 | 61 | 3.20 | 756 | 4.61 |
| 02 | 128 | 5.35 | 1226 | 6.54 |
| 03 | 102 | 4.2 | 659 | 4.01 |
| 04 | 69 | 3.67 | 192 | 2.36 |
| 05 | 33 | 1.38 | 287 | 2.53 |
| 06 | 130 | 2.51 | 229 | 2.35 |
| 07 | 45 | 2.1 | 538 | 3.58 |
| 08 | 51 | 1.58 | 223 | 2.21 |
| 09 | 25 | 2.08 | 651 | 3.67 |
| 10 | 92 | 2.54 | 854 | 5.30 |
| 11 | 77 | 3.1 | - | - |
| 12 | 41 | 1.33 | - | - |
| 13 | 23 | 2.33 | - | - |
| 14 | 87 | 3.28 | - | - |
| 15 | 44 | 2.39 | - | - |
| 16 | 45 | 3.03 | - | - |
| 17 | 36 | 1.56 | - | - |
| 18 | 64 | 3.12 | - | - |
| 19 | 61 | 3.20 | - | - |
| 20 | 67 | 3.40 | - | - |
| Mean±SEM | 64 ± 6.92 | 2.7 ± 0.23 | 561 ± 75.2 | 3.72 ± 0.32 |

